# Flexible and efficient handling of nanopore sequencing signal data with *slow5tools*

**DOI:** 10.1101/2022.06.19.496732

**Authors:** Hiruna Samarakoon, James M. Ferguson, Sasha P. Jenner, Timothy G. Amos, Sri Parameswaran, Hasindu Gamaarachchi, Ira W. Deveson

## Abstract

**Background:** Nanopore sequencing is an emerging technology that is being rapidly adopted in research and clinical genomics. We recently developed SLOW5, a new file format for storage and analysis of raw data from nanopore sequencing experiments. SLOW5 is a community-centric, open source format that offers considerable performance benefits over the existing nanopore data format, known as FAST5. Here we introduce *slow5tools*, a simple, intuitive toolkit for handling nanopore raw signal data in SLOW5 format.

**Results:** *Slow5tools* enables lossless FAST5-to-SLOW5 and SLOW5-to-FAST5 data conversion, and a range of tools for structuring, indexing, viewing and querying SLOW5 files. *Slow5tools* uses multi-threading, multi-processing and other engineering strategies to achieve fast data conversion and manipulation, including live FAST5-to-SLOW5 conversion during sequencing. We outline a series of examples and benchmarking experiments to illustrate *slow5tools* usage, and describe the engineering principles underpinning its high performance.

**Conclusion:** *Slow5tools* is an essential toolkit for handling nanopore signal data, which was developed to support adoption of SLOW5 by the nanopore community. *Slow5tools* is written in C/C++ with minimal dependencies and is freely available as an open-source program under an MIT licence: https://github.com/hasindu2008/slow5tools.

## INTRODUCTION

Nanopore sequencing analyses native DNA or RNA molecules with no upper limit on read length^1^. Popular devices from Oxford Nanopore Technologies (ONT) measure the displacement of ionic current as a DNA/RNA strand passes through a protein pore, recording time-series signal data in a format known as ‘FAST5’. This signal data is typically ‘base-called’ into sequence reads and is further used during a wide range of downstream analyses^2–8^.

ONT’s FAST5 data format suffers from several inherent limitations, which we have articulated previously^9^. FAST5 file sizes are inflated by inefficient space allocation and metadata redundancy. Its dependence on Hierarchical Data Format (HDF) makes FAST5 unsuitable for efficient parallel access by multiple CPU threads. FAST5 files can only be read/written by the *hdf5lib* software library, which reduces usability and can make even relatively simple operations difficult and expensive.

We recently developed a new file format named SLOW5, which is designed to overcome the above limitations in FAST5^9^. In its compressed binary form (BLOW5), the new format is ∼20-80% smaller than FAST5 and permits efficient parallel access by multiple CPU threads. This enables order-of-magnitude improvements in the speed and cost of signal data analysis on common high-performance computing (HPC) and cloud systems^9^. SLOW5 is well documented and fully open source. Data reading/writing is managed by an intuitive software API that is compatible with a wide range of system architectures. Overall, SLOW5 offers many benefits to the ONT user community.

To accompany the SLOW5 data format, we have developed a suite of tools for creating, handling and interacting with SLOW5/BLOW5 files. *Slow5tools* includes all utilities required for novice and advanced users to integrate SLOW5 with their workflows, and solves existing challenges in the management and analysis of ONT raw signal data. *Slow5tools* is implemented in C/C++ and utilises multi-threading, multi-processing and other software engineering strategies to achieve fast and efficient performance. This article provides a series of examples that serve to articulate the design, usage and performance of *slow5tools*.

## RESULTS

### Overview of slow5tools

*Slow5tools* is a modular toolkit for working with signal data from ONT experiments. *Slow5tools* is written in C/C++ and uses two core libraries, *slow5lib* and *hdf5lib*, for reading/writing SLOW5/BLOW5 and FAST5 files, respectively (see **Methods**; **Fig. S1a**). The software has the following basic usage, where ‘command’ specifies the operation to be performed:

~~~
# generic usage for slow5tools
slow5tools command input.slow5 -o output.slow5
~~~

The various commands currently available in *slowt5tools* are summarised in **Figure 1** and **Table S1**. Different commands within *slow5tools* employ various engineering strategies to achieve optimum performance. These strategies are articulated below, along with relevant benchmarking results.

**Figure 1.**
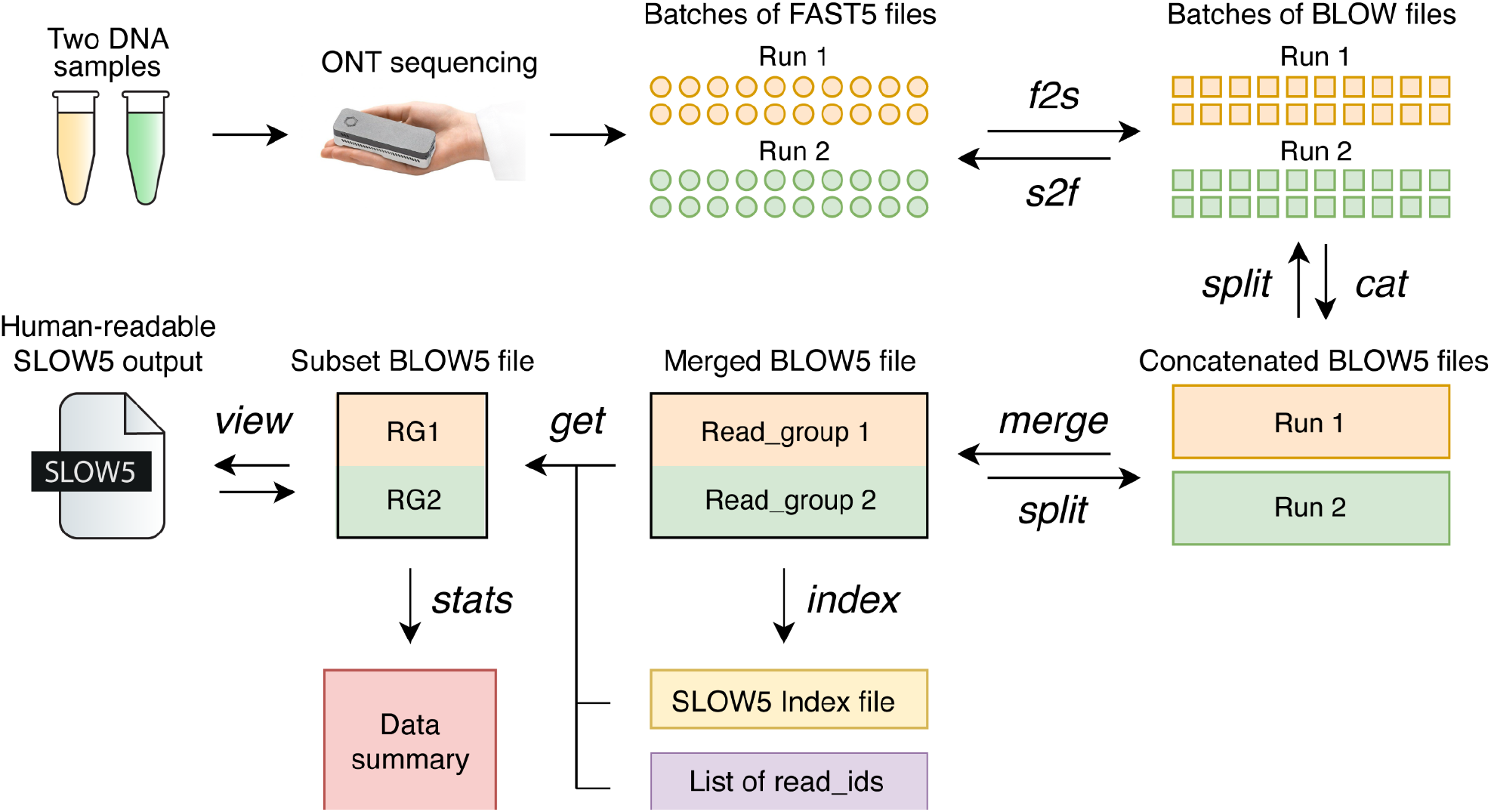
Schematic overview of nanopore raw signal data management with *slow5tools*. The software permits lossless data conversion from FAST5-to-SLOW5 (*f2s* command) or SLOW5-to-FAST5 (*s2f* command). *Slow5tools* executes a range of common operations for data management, including combining multiple SLOW5/BLOW5 datasets (*merge* or *cat* commands) or splitting a dataset into multiple parts (*split* command). The *index* command creates an index file for a SLOW5/BLOW5 file, which then enables a specific record(s) to be quickly fetched from the file using the *get* command. The *view* command is used to view the contents of a binary BLOW5 file/s in human-readable SLOW5 format, convert datasets between SLOW5 and BLOW5, or between different BLOW5 compression types. Finally, *slow5tools* can check the integrity of a SLOW5/BLOW5 file (*quickcheck*) or generate simple summary statistics describing the dataset (*stats*).

### Fast, lossless data conversion

Because ONT devices currently write data in FAST5 format, data conversion to SLOW5/BLOW5 format will be the first step in most workflows. FAST5-to-SLOW5 conversion can be performed with *slow5tools f2s*, as follows:

~~~
# convert a single FAST5 file into SLOW5 ASCII format
slow5tools f2s file.fast5 -o file.slow5
# convert a directory of FAST5 files into binary BLOW5 files
slow5tools f2s fast5_dir -d blow5_dir
~~~

Data conversion is lossless by default, and the user may convert their data back to the original FAST5 format using *slow5tools s2f*, as follows:

~~~
# convert a directory of BLOW5 files to FAST5
slow5tools s2f blow5_dir -d fast5_dir
~~~

It is not possible to read/write FAST5 files using parallel CPU threads, due to their dependence on the *hdf5lib* library, which serialises any thread-based input/output (I/O) operations^9^. Therefore, *f2s/s2f* data conversion is instead parallelised through a multi-processing strategy illustrated in **Fig. S1b** (see **Methods**). To evaluate this solution, we measured runtimes for data conversion on a typical ONT sequencing dataset (∼9M reads; **Table S2**), as executed with various numbers of processes on three machines with different disk systems (SDD, HDD, Distributed; **Table S3**). Data conversion rates scaled with the number of processes for both *f2s* and *s2f*, and the observed trends were similar to the ideal curves, indicating highly efficient utilisation of parallel compute resources, with the exception of System 2 (**Figure 2a,b**). Data conversion on System 2 showed sub-optimal parallelism for two main reasons: (*i*) System 2 has a traditional HDD disk system that throttles when the number of I/O requests is high; (*ii*) using processes instead of threads makes these I/O requests completely separated and thus the associated overhead is high. This experiment demonstrates that multi-processing is a viable solution for parallelisation of FAST5/SLOW5 data conversion, in which multi-threading is not permitted, although this approach has limitations that will impact performance on some systems.

**Figure 2.**
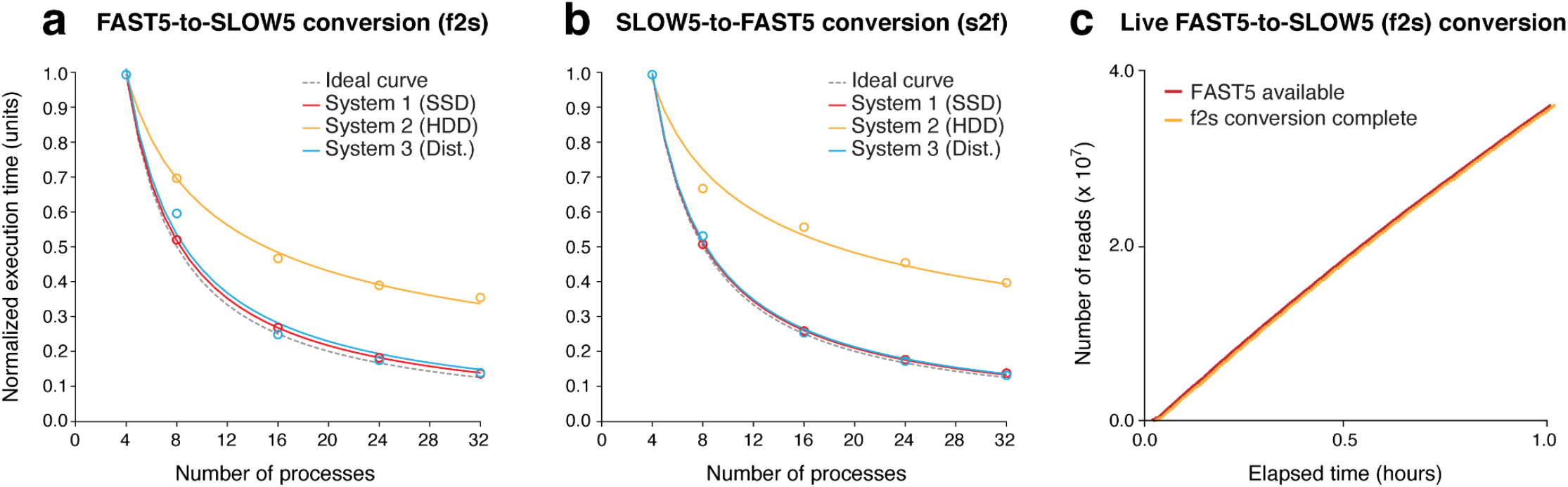
Efficient data conversion with *slow5tools f2s* and *s2f* commands. (**a**,**b**) Normalised execution time for conversion of a typical ONT dataset (∼9M reads) from FAST5-to-BLOW5 format (**a**), or BLOW5-to-FAST5 format (**b**) relative to the number of parallel processes used by *slow5tools*. Data conversion was evaluated on three separate machines with different disk systems (SSD, HDD, Distributed). Ideal curves (dashed line) model the expected performance under optimal parallelism. (**c**) The rate of FAST5 data production (red) and FAST5-to-BLOW5 data conversion (orange) with *realf2s*.*sh* on a PromethION P48 device running at maximum sequencing load.

With the speeds achieved by *f2s* above, live FAST5-to-SLOW5 data conversion during a sequencing experiment is also possible. To manage this process, we developed an accompanying script (*realf2s*.*sh*) that detects freshly created FAST5 files during sequencing, converts them to BLOW5, and checks the integrity of the data before (optionally) deleting the FAST5s (see **Methods**). We tested this feature on ONT’s PromethION P48 device, using the onboard computer for live data conversion (**Table S3**), and found that *f2s* can easily match the pace of data production at maximum sequencing load (i.e. 48 flow cells running in parallel; **Figure 2c**). Indeed, the average delay between the completion of an individual FAST5 file containing 4000 reads and its conversion to BLOW5 format was just ∼20 seconds. Given the smaller size of BLOW5 vs FAST5 files (typically ∼50% smaller), live *f2s* conversion increases the number of sequencing runs that can be performed in parallel before reaching the maximum storage capacity on an ONT sequencing device. We also anticipate that the use of live-converted BLOW5 format will also improve the capacity of live base-calling that is achievable on ONT sequencing devices in the future.

### Merging and splitting datasets

The *slow5tools merge* command can also be used to combine multiple batches of reads or data from different sequencing runs into a single BLOW5 file, for example when aggregating data across replicate experiments. If data is merged from different runs/flow-cells, each will be assigned a separate *read_group*, with which all its constituent reads are tagged in the merged file. Merging can be performed as follows:

~~~
# merge all BLOW5 files in one or more directories into a single BLOW5 file
slow5tools merge blow5_dir1 blow5_dir2 -o file.blow5
~~~

Data merging is accelerated via an interleaved multi-threading strategy illustrated in **Fig. S1c** (see **Methods**), which is more resource-efficient than the multi-processing strategy described above. Evaluating this solution on the same machines as above, we observed near-optimal parallelism, with data merging times being reduced in proportion to the number of CPU threads used (**Figure 3a, Fig. S2a**).

**Figure 3.**
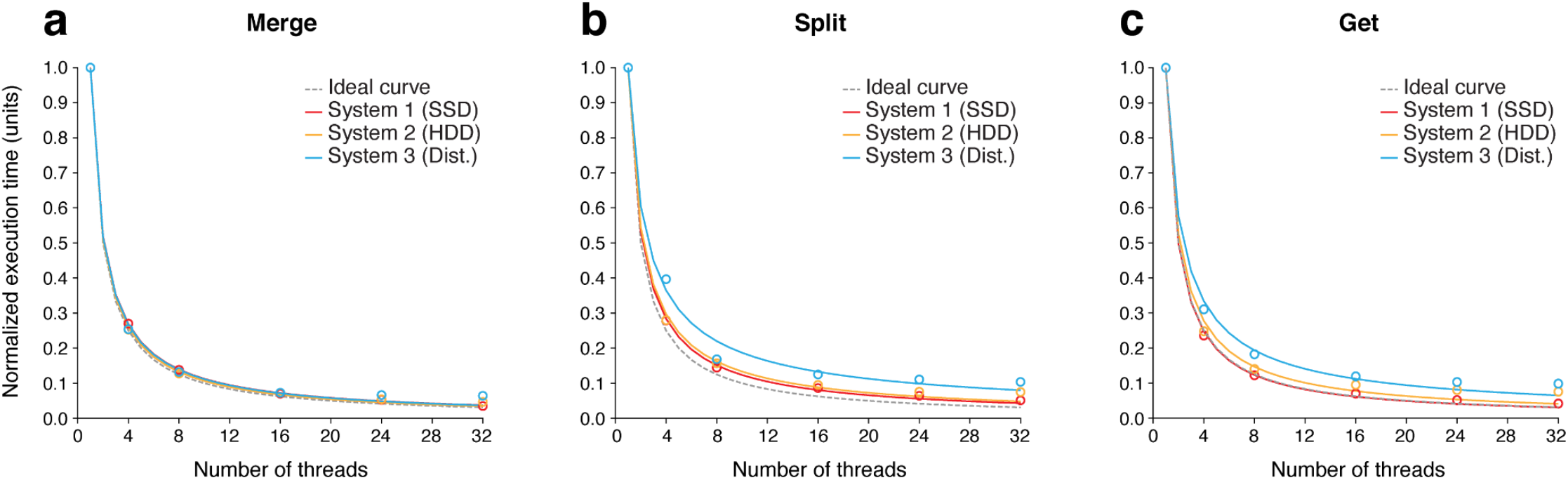
Merging, splitting and querying BLOW5 files. (**a**) Normalised execution time for merging all individual batches of reads produced by MinKNOW into a single BLOW5 file for a typical ONT dataset (∼9M reads), relative to the number of parallel CPU threads used by *slow5tools*. Data merging was evaluated on three separate machines with different disk systems (SSD, HDD, Distributed). Ideal curves (dashed line) model the expected performance under optimal parallelism. (**b**) The same comparison for splitting a typical BLOW5 file into separate files for each read_group with *slow5tools split*. (**c**) The same comparison for retrieving a set of records from a BLOW5 file based on their read_ids with *slow5tools get*.

The *slow5tools cat* command provides a lightweight alternative to *merge*, which can be used to quickly combine batches of reads without assigning read_groups. Similar to the Unix ‘cat’ command, this tool is intended for concatenating the small batches of reads produced by ONT’s MinKNOW software into a single BLOW5 file for a given sequencing run, and does so faster than possible with *merge* (**Fig. S2b**) *Cat* does not permit the user to combine files with different metadata (e.g. reads from different flow cells) in their headers.

The user may also wish to split a dataset into multiple separate files. This can be achieved using the *split* command, which offers the user a choice to direct reads from each *read_group* to a single output file, or alternatively split the input into a specific number of output files or a specific number of reads per file:

~~~
# split a BLOW5 file into separate BLOW5 files based on read groups
slow5tools split file.blow5 -d blow5_dir -g
# split a BLOW5 file (single read group) into separate BLOW5 files such that there
are 4000 reads per file
slow5tools split file.blow5 -d blow5_dir -r 4000
# split a BLOW5 file (single read group) into 100 separate BLOW5 files
slow5tools split file.blow5 -d blow5_dir -f 100
~~~

When splitting by read_group, *split* is accelerated via a multi-threading strategy similar to *merge* (see **Methods**), and also exhibits near-optimal parallelism on different machines (**Figure 3b**). Together, the *merge, cat* and *split* tools provide the user an efficient, flexible framework for reorganising ONT raw signal datasets into the most convenient structure for any use case.

### Indexing SLOW5/BLOW5 files

Efficient access to a SLOW5/BLOW5 file is facilitated by an accompanying binary index file that specifies the position of each read (in bytes) within the main file (see **Methods**). This file should be created using the *slow5tools index* command:

~~~
# index a SLOW5/BLOW5 file
slow5tools index file.blow5
~~~

Creating an index file for a BLOW5 file involves decompressing and parsing each record to get the position of each read_id. Decompressing and parsing an entire record is time consuming. However, the zlib compression algorithm, used by default in *slow5tools*, supports partial decompression. *Index* is hence accelerated through a partial decompression strategy, where only the read_id is decompressed for a given record. This method enjoys substantial performance benefits during index creation, compared to complete decompression (**Fig. S2c**).

### Viewing and querying SLOW5/BLOW5 files

SLOW5 datasets can be stored in human-readable ASCII SLOW5 format, or the more compact binary BLOW5 format, which is preferable in most scenarios. The *view* tool allows the user to view the contents of a BLOW5 file or convert between SLOW5/BLOW5:

~~~
# to view a BLOW5 file in SLOW5 ASCII on standard output
slow5tools view file.blow5
# Convert a BLOW5 file into SLOW5 ASCII
slow5tools view file.blow5 -o file.slow5
# convert a SLOW5 file to BLOW5 (default compression)
slow5tools view file.slow5 -o file.blow5
~~~

The *view* command can also be used to specify different compression methods available for BLOW5 format, with various combinations available across two compression layers (see **Methods**).

Rather than viewing the whole file, many use-cases require retrieval of a specific record/s from a dataset, based on their read_ids. This can be performed with the *slow5tools get* command, which enables efficient random access on BLOW5 files, as follows:

~~~
# extract records from a SLOW5/BLOW5 file corresponding to given read ids
slow5tools get file.blow5 readid1 readid2 -o output.slow5
~~~

*Get* employs a similar multi-threading strategy as above, and exhibits optimal parallelism on all systems tested (**Figure 3c**). Together these tools allow users to efficiently view and query datasets in SLOW5/BLOW5 format.

### Other tools

In addition to the commands outlined above, *slow5tools* also currently includes two tools for checking the integrity and/or specifications of a dataset in SLOW5/BLOW5 format. The *slow5tools stats* command generates simple summary statistics including the read count, number of read_groups, etc. A lightweight alternative, *quickcheck* simply checks a SLOW5/BLOW5 file to make sure it is not malformed. The modular architecture of *slow5tools* will allow new tools to be readily added as new use cases arise. Similarly, the *slow5tools* command structure enables multiple operations to be combined in efficient one-line BASH commands familiar to the bioinformatics community (https://hasindu2008.github.io/slow5tools/oneliners.html).

## DISCUSSION

SLOW5 format was developed as an open-source, community-centric file format that addresses several inherent design-flaws in ONT’s FAST5 format, on which the nanopore community was previously dependent^9^. Performance, compatibility, usability and transparency are central principles of the ongoing SLOW5 initiative.

In addition to the *slow5lib* and *pyslow5* APIs for reading and writing SLOW5 files, *slow5tools* is critical for the usability of SLOW5 format. With ONT devices currently limited to write data in FAST5 format, the *slow5tools f2s* utility for FAST5-to-SLOW5 conversion will be the first step in any workflow that integrates SLOW5. We also provide a method for live data conversion during sequencing, which easily keeps pace with a PromethION P48 device at maximum load. This allows the user to obtain their data in analysis-ready BLOW5 format with no additional cost to the overall workflow time, or to more efficiently move raw data off the sequencing machine for real-time analysis^10^. This also provides a base for future implementation of real-time basecalling with SLOW5 input, which we expect will enjoy significant performance benefits over FAST5.

Following data conversion, *slow5tools* provides a fast and flexible framework for managing and interacting with SLOW5/BLOW5 files. We recommend the use of binary BLOW5 for almost all applications, because it is more compact and efficient than ASCII SLOW5. BLOW5 supports multiple compression methods, or no compression, at the user’s discretion. Zlib was chosen as the default compression method due to its near-universal compatibility, however svb+zstd is more compact and provides superior file-reading performance. In some cases, such as when writing temporary BLOW5 files during an analysis, no compression may also be preferable. SLOW5 was designed to allow future compression strategies to be easily incorporated, and we anticipate domain-specific compression algorithms will deliver future improvements.

*Slow5tools* is intended to facilitate adoption of SLOW5 among ONT users. Given the performance improvements of SLOW5 over FAST5^9^, this will be of significant benefit to the ONT community. Equally important, *slow5tools* greatly simplifies the basic processes involved in structuring, querying and interacting with nanopore raw signal data files, which are at the heart of all data management and analysis workflows. We provide *slow5tools* as a free, open-source software package: https://github.com/hasindu2008/slow5tools.

## METHODS

### Basic architecture/implementation of slow5tools

Slowtools is written in C/C++ and links with two file format specific libraries. *Hdf5lib* and *slow5lib* are used to read and write files in FAST5 and SLOW5/BLOW5 formats, respectively (**Fig. S1a**). There are different FAST5 versions (v2.2 is latest at time of submission), and all current/previous versions can be converted to SLOW5 format using *slow5tools*. When converting back to FAST5 format, *slow5tools* conforms to the latest FAST5 version (currently v2.2). *Slow5tools* will provide support for future FAST5 versions. SLOW5 file format specification (https://hasindu2008.github.io/slow5specs/) is maintained separately from *slow5lib* and *slow5tools*. Backwards compatibility will be ensured in future SLOW5 versions and a compatibility table is maintained here: https://hasindu2008.github.io/slow5tools/compatibility.html

### Slow5tools multi-processing strategy

All *slow5tools* commands are optimised to run efficiently on modern computer CPUs. The key optimization is to exploit parallelisation. Both the *f2s* and *s2f* commands require *hdf5lib* to read and write FAST5 files, respectively. *Hdf5lib* internally uses global locks to serialise parallel thread-based I/O operations. Therefore, to parallelise the conversion from FAST5 to SLOW5 and vice versa, we employ a multi-processing strategy (**Fig. S1b**). In this approach, each process keeps a separate memory instance of the library. Firstly, the program recursively searches the input directories to list the files to be converted based on the file extension (e.g. *f2s* lists all files with .fast5 extensions). Then, using the C function fork(), a pool of processes is created. The user can determine the number of processes to spawn (default is 8). Next, the number of files are equally distributed among the processes. Inside a process, an input file is opened, an output file is created, the content of the input file is written to the output file as per the output file format and lastly, both files are closed. Then the process iterates to the next alloted input file, if any. When all the processes terminate all the files are converted successfully.

### Slow5tools multi-threading strategy

To exploit parallelisation in the rest of the programs (where FAST5 files are not involved), a multithreading model is adopted (**Fig. S1c**), which uses less resources than a similar multiprocessing model. This is possible as these programs only depend on slow5lib that supports C/C++ POSIX threads. A SLOW5 record has a finite set of primary attributes - read_id, read_group, digitisation, offset, range, sampling rate, len_raw_signal, and the raw signal. It can optionally have auxiliary attributes. By default, the raw signal is compressed using the stream variable byte (svb) encoding algorithm and the whole record is compressed using standard zlib compression (see below). When reading a SLOW5 file, first the compressed read should be read from the storage disk, decompressed and then parsed into structuresin the memory. These steps can exploit parallelism to speed up the I/O operations. That is, a single threaded application (main program) fetches a batch of compressed records (using *slow5_get_next_mem()*) from the file on the disk and spawns a set of worker threads to decompress and parse (using *slow5_rec_depress_parse()*) them. Later, the necessary processing on the records is also carried out on the threads. The output records are written to a buffer (using *slow5_rec_to_mem()*) that is accessible by the main program’s thread before the worker threads exit. Eventually, after returning to the main thread (after worker threads are joined by the main thread), the content in the buffer is written to the output stream. This workflow is carried out until the end of the file is reached. The number of records to fetch at a time is determined by the batch size parameter of the programs. The user can also set the number of threads to match the available compute resources. This method ensures the order of the records of the input is maintained in the output.

### Single thread implementations

*Slow5tools stats* gives a summary of the slow5 file. In addition to the details about the primary and auxiliary attributes it counts the number of reads in the file. Since no processing is done on the records, a single threaded program that continuously calls *slow5_get_next_mem()* on the file is the optimal method to count the records. However, the buffer size of the standard C function *fread()* determines how many bytes are fetched from the file through system calls. This affects the performance of *slow5_get_next_mem()* and thus *slow5tools*. Hence, the buffer size was set to a larger buffer size of 128k bytes using *setvbuf()* in *stdio*. To split a SLOW5/BLOW5 file into a given number of reads or files, *split* also uses a single threaded implementation unless the user sets a different output format from the input format. That is because this requires no processing on a record but just writing as it is to an output file.

### BLOW5 compression strategies

There are two compression layers applied on a SLOW5 record. By default, BLOW5 format has zlib compression and svb-zd encoding [zlib, svb-zd] on the entire record and on the signal respectively. *Slow5tools view* can be used to create BLOW5 files with different compression combinations. View also adopts the multithreading model explained above (**Fig. S1c**). For example, to convert a BLOW5 record from [zlib, svb-zd] to [zlib, none] (i.e. no signal encoding), the record should be first decompressed using zlib followed by a svb-zd decoding of the signal. Then the records are compressed back using zlib. On average, this process is ∼6X times slower than the conversion from [zlib, svb-zd] to [zstd, svb-zd], because zlib compression is slower than zstd compression. Despite this disparity, zlib compression is used as the default BLOW5 record compression method for the sake of compatibility. Zstd library is not yet widely available by default, specially on academic HPC servers and the user may face associated dependency and version issues. However, we advise the users to use zstd compression where this is possible on their specific system.

### Indexing

*Slow5tools index* creates a binary index file that stores the read_ids and the byte offset of the records. Loading the index to memory allows bioinformatics programs to quickly fetch the necessary records from a SLOW5/BOW5 file in a random access pattern. Creating an index file from a BLOW5 file involves decompressing and parsing each record to get the read_ids. Decompressing and parsing an entire record just to fetch the read_id is time consuming. The default SLOW5 record compression method, zlib, supports partial decompression. This enables quick access to the read_id by only decompressing up to the read_id of a record. The other compression method, zstd, does not support partial decompression. Therefore, *index* first determines which compression is used, and then, either partially or completely decompresses the BLOW5 record to fetch the read_id. Using this partial decompression method, indexing is ∼15 times faster than with the default zlib decompression and, despite the fact that zlib is slower than zstd, indexing is ∼3 times faster with zlib partial decompression than zstd (which must fully decompress each record) (**Fig. S2c**).

### Live data conversion

Live FAST5-to-SLOW5 data conversion can be carried out using the *realf2s*.*sh* script provided under the *slow5tools* repository at https://github.com/hasindu2008/slow5tools/tree/master/scripts/realtime-f2s. This is a BASH script that takes the sequencing data directory as an argument (e.g., /data/sample_id on a standard PromethION architecture). The script continuously monitors this specified directory for newly created FAST5 files using the *inotifywait* tool available under Linux’s inotify interface. As soon as a newly created FAST5 file is detected (the default in MinKNOW is such that one FAST5 file contains 4000 reads), the *slow5tools f2s* command is invoked unless the maximum number of processes is already reached. If the maximum number of processes has been reached, the newly detected FAST5 files are queued until the ongoing conversions complete. This strategy prevents the system being overloaded if hundreds of files are generated by MinKNOW at once.

To test the capacity of real-time data conversion on a PromethION computer tower, we simulated 48 parallel sequencing runs using the playback feature in MinKNOW. The bulk-FAST5 file for MinKNOW playback was saved during sequencing of a typical human genome sample (∼10-20kb reads, R9.4.1 flow cell, LSK110 kit). The simulation was performed by modifying the relavent MinKNOW TOML file (TOML file named *sequencing_PRO002_DNA*.*toml* under */opt/ont/minknow/conf/package/sequencing* was modified to include the “*simulation = /path/to/bulk/fast5”* field under “*[custom_settings]”* section)) to point to this bulk-FAST5 file. The simulation was run for a period of 1 hour with live basecalling disabled. The rate of data generation is highest during the first hour of a sequencing run; therefore this simulation emulates the maximum load one could expect to encounter. Live basecalling was disabled during the simulation so that completed FAST5 files are released at the sequencing rate - this would otherwise be tapered by MinKNOW because live basecalling cannot keep up with the sequencing rate. For this experiment, the maximum number of processes was capped at 56 (half of the CPU threads available on the PromethION compute-node), thus limiting the maximum number of FAST5 files simultaneously converted to be 56.

## DATA & CODE AVAILABILITY

SLOW5 format and all associated software are free and open source. Datasets used in benchmarking experiments are described in **Table S2** and are available on NCBI Sequence Read Archive at Bioproject PRJNA744329.

*Slow5tools* can be accessed at: https://hasindu2008.github.io/slow5tools/

*Slow5lib & pyslow5* can be accessed at: https://hasindu2008.github.io/slow5lib/

SLOW5 format specification documents can be accessed at: https://hasindu2008.github.io/slow5specs

## ACKNOWLEDGEMENTS

We thank our colleague Derrick Lin for providing excellent technical support and freedom to use the institute’s HPC system in some quite exotic ways. Resources from the Australian National Computational Infrastructure (NCI) were used during benchmarking experiments. We acknowledge the following funding support: Australian Medical Research Futures Fund Investigator Grant MRF1173594 (to I.W.D.) and philanthropic support from The Kinghorn Foundation (to I.W.D.).

## DECLARATIONS

I.W.D. manages a fee-for-service sequencing facility at the Garvan Institute of Medical Research that is a customer of Oxford Nanopore Technologies but has no further financial relationship. H.G. & J.M.F. have previously received travel and accommodation expenses to speak at Oxford Nanopore Technologies conferences. The authors declare no other competing financial or non-financial interests.

## CONTRIBUTIONS

All authors (H.S., J.M.F., S.P.J., T.G.A., S.P., H.G., & I.W.D.) contributed to the conception, design and testing of *slow5tools*. H.S., J.M.F., S.P.J., & H.G. implemented core *slow5tools* software. H.S. & H.G. performed benchmarking experiments. H.S. & I.W.D. prepared the figures. H.S., H.G. & I.W.D prepared the manuscript, with support from all authors.

