## Supplementary_Materials for "Flexible and efficient handling of nanopore sequencing signal data with *slow5tools*"

**Table S1. Summary of current commands in *slow5tools*.**

| Command | Function |
|---|---|
| f2s | Convert FAST5 files to SLOW5/BLOW5 format. |
| s2f | Convert SLOW5/BLOW5 files to FAST5 format. |
| merge | Merge multiple SLOW5/BLOW5 files to a single file. |
| cat | Quickly concatenate SLOW5/BLOW5 files of the same read group to a single file. |
| split | Split a single SLOW5/BLOW5 file into multiple separate files. |
| view | View the contents of a SLOW5/BLOW5 file or convert between different SLOW5/BLOW5 formats and compressions. |
| get | Retrieve records for specified read IDs from a SLOW5/BLOW5 file. |
| stats | Prints summary statistics describing a SLOW5/BLOW5 file. |
| quickcheck | Quickly checks if a SLOW5/BLOW5 file is intact. |
| help | Prints the help message. |
| version | Prints the version. |

**Table S2. Data specifications.**

| Dataset | Platform | Pore | Reads | Total seq. | Size FAST5 | Size BLOW5 (zlib+svb-zd) | Size BLOW5 (zstd+svb-zd) |
|---|---|---|---|---|---|---|---|
| ~30X human genome (NA12878) | PromethION | R9.4.1 | 9,083,052 | 93.4 Gbases | 1.3 TB | 0.74TB | 0.71TB |

**Table S3. Computer specifications.**

| System | Name | CPU (No. of cores) | RAM | File system | Disk system | OS |
|---|---|---|---|---|---|---|
| 1 | HPC with SSD | 2 x Intel Xeon Gold 6140M (72) | 3000 GB | ext4 | 6x4TB NVME SSD drives with RAID0 configuration | CentOS 7.9.2009 |
| 2 | HPC with HDD RAID | 2 × Intel Xeon Gold 6154 (36) | 384 GB | ext4 | 12×10TB HDD drives with RAID6 configuration | Ubuntu 18.04.5 LTS |
| 3 | NCI-Gadi CPU node with distributed lustre file system | 2 x Intel Xeon Platinum 8274 (48) | 192 GB | lustre | 7200 4TB disks in 120 NetApp disk arrays | CentOS 8.3.2011 |
| 4 | PromethION P48 compute node | 2x Intel Xeon Platinum 8180 (112) | 384 GB | ext4 | 8x8TB SSD drives with RAID0 configuration | Ubuntu 20.04.4 LTS |

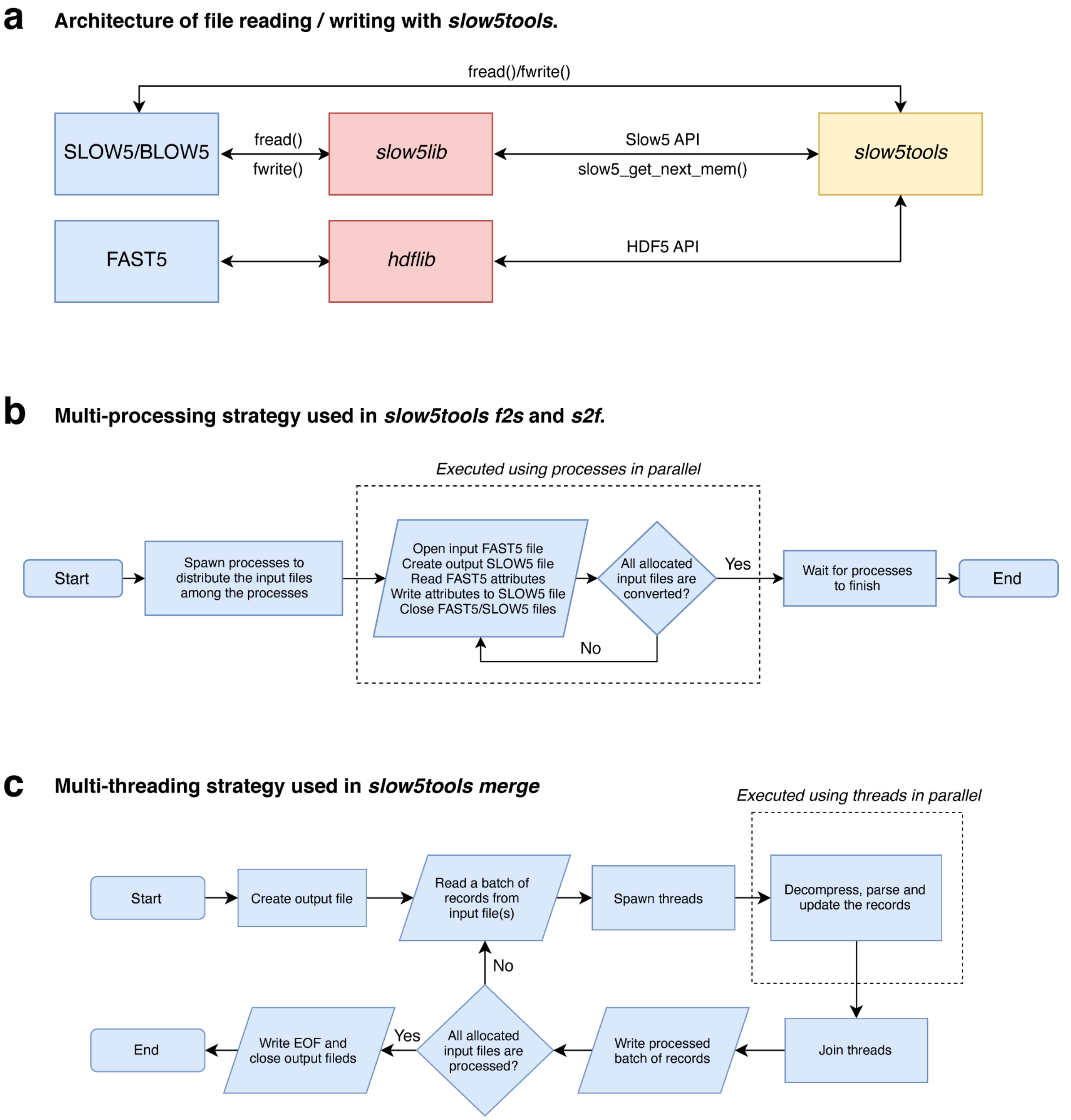


**Fig. S1. *Slow5tools* design principles and optimisations.** (**a**) *Slow5tools* uses *slow5lib* and *hdf5lib* to read and write files in SLOW5/BLOW5 and FAST5 formats, respectively. *Slow5tools* uses the library API functions except in the case of *cat* which directly reads and writes using *fread* and *fwrite* for better performance. *Slow5lib* provides fine grained API functions such as *slow5_get_next_mem()* to implement parallel execution. (**b**) *F2s* and *s2f* are implemented in an embarrassingly parallel manner. A pool of processes is created at the beginning. Each process converts FAST5 files to SLOW5/BLOW5 format independently. (**c**) The underlying multithreading strategy adopted by *merge, split, get,* and *view* is very similar. A batch of raw records are fetched, processed, and written to the output iteratively. The processing is executed using threads in parallel. In the *merge* implementation, the three stages are synchronised (not shown).

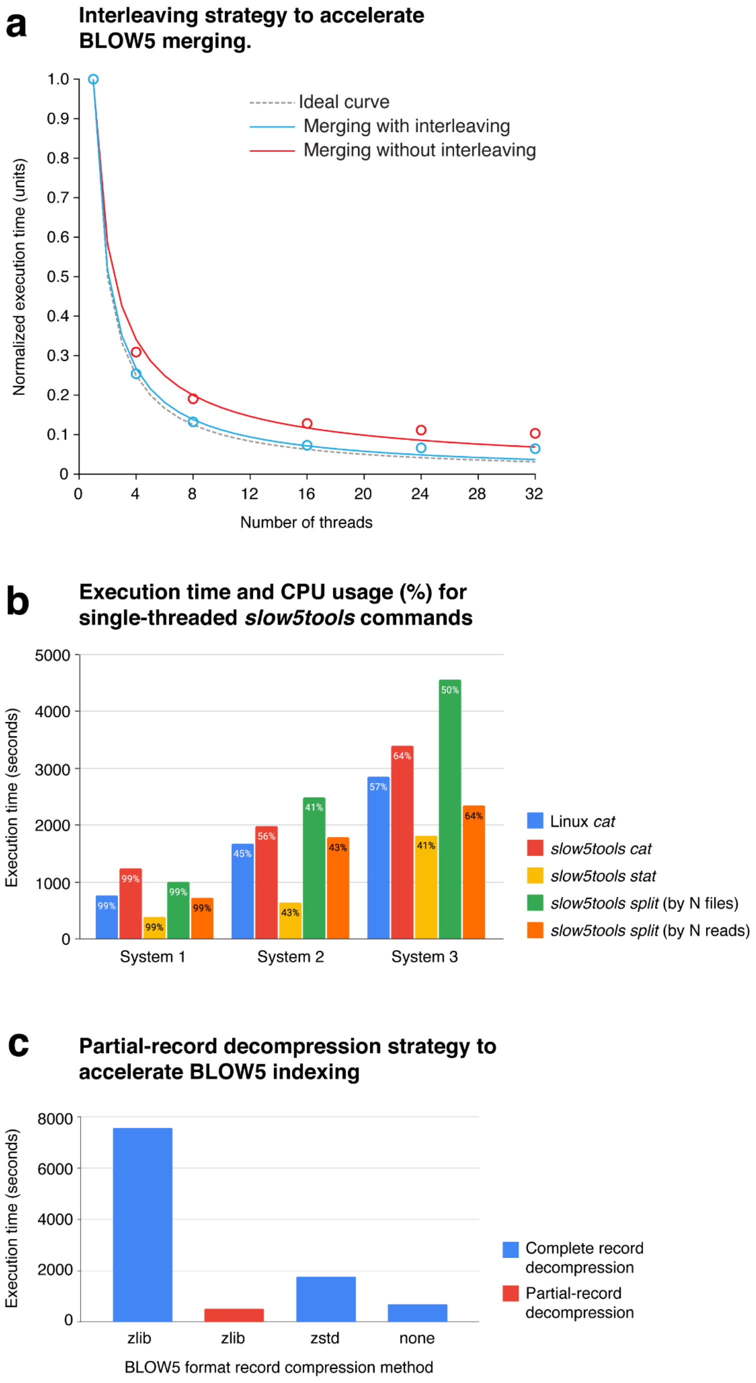


**Fig. S2. *Slow5tools* acceleration strategies.** (**a**) *Merge* has three stages of execution running on a loop; reading a batch of reads, processing and writing them to the output. When muti-threading was used for processing, all three stages recorded equal timings making it possible to create a pipelined execution; I/O became the bottleneck that could be overlapped with a stage from the previous iteration. This interleaved strategy improves the efficiency of merging, and showed a trend close to the trend of the expected performance under optimal parallelism. (**b**) *Slow5tools cat* command is very similar to Unix *cat* command with more sanity checks that are specific to SLOW5 format. Both *Slow5tools cat* and Unix *cat* have similar performance across all the systems. As expected, the *slow5tools stats* command takes the minimum execution time as it does not process records, but counts them. All the programs showed almost 100% CPU utilisation on System 1 that uses an SSD disk system. This guarantees that the increased execution timings observed on Systems 2 and 3 were solely because of the slower I/O performance compared to SSD. (**c**) Indexing involves decompressing, parsing and fetching the *read_id* from the SLOW5 records. Partially decompressing only up to the *read_id* (only possible with zlib) instead of a full record decompression dramatically improved the execution time.
